# Laboratory cultivation of acidophilic nanoorganisms. Physiological and bioinformatic dissection of a stable laboratory co-culture

**DOI:** 10.1101/103150

**Authors:** Susanne Krause, Andreas Bremges, Philipp C. Münch, Alice C. McHardy, Johannes Gescher

## Abstract

This study describes the laboratory cultivation of ARMAN (Archaeal Richmond Mine Acidophilic Nanoorganisms). After 2.5 years of successive transfers in an anoxic medium containing ferric sulfate as an electron acceptor, a consortium was attained that is comprised of two members of the order *Thermoplasmatales*, a member of a proposed ARMAN group, as well as a fungus. The 16S rRNA of one archaeon is only 91.6% identical to *Thermogymnomonas acidicola* as most closely related isolate. Hence, this organism is the first member of a new genus. The enrichment culture is dominated by this microorganism and the ARMAN. The third archaeon in the community seems to be present in minor quantities and has a 100% 16S rRNA identity to the recently isolated *Cuniculiplasma divulgatum*. The enriched ARMAN species is most probably incapable of sugar metabolism because the key genes for sugar catabolism and anabolism could not be identified in the metagenome. Metatranscriptomic analysis suggests that the TCA cycle funneled with amino acids is the main metabolic pathway used by the archaea of the community. Microscopic analysis revealed that growth of the ARMAN is supported by the formation of cell aggregates. These might enable cross feeding by other community members to the ARMAN.

## Introduction

The question of the smallest autonomous living cell is an intensely discussed topic in microbiology. In 1999, the Steering Group for the Workshop on Size Limits of Very Small Microorganisms defined the lower theoretical limit for an autonomous living cell inhabiting all necessary components to 0.008-0.014 µm^3^^1^. Early findings of ultra-small microorganisms, e.g. “ultramicrobacteria” in sea-water^2^ and “dwarf cells” or “nanobacteria” in soils^3,4^, and the question of the potential medical relevance^5^ of some of these cells, were the starting point and motivation for a new research direction. Nevertheless, the existence of some of these nanobacteria were questioned later for being microcrystalline apatite rather than living cells^6^ or could at least be starvation forms^7^. Still, there are several organisms with cell sizes close to or even below the mentioned theoretical limit that were identified in different habitats in the last century.

Filtration of water samples with 0.2 or 0.1 µm filters and subsequent sequencing of the 16S rRNA genes of the organisms has revealed that the enriched small or ultra-small microbes belong to candidate divisions that are widespread in nature but lack any cultured representative^8,9^. Luef *et al.* (2015) even hypothesized that it might be a general characteristic of previously uncultivable bacterial phyla that the organisms are small^9^. Thus far, isolated ultramicrobacteria belong to different phyla including *Actinobacteria*^10^, *Bacteroidetes*^11^ or *Alphaproteobacteria*^12,13^. They have cell volumes between <0.1 µm³ and <0.004 µm³, and their energy metabolism is described as chemoorganotrophic, heterotrophic or facultatively parasitic. Nevertheless, detailed information about optimal growth conditions, ecological role and corresponding genomic or transcriptomic data are often not available.

*Nanoarchaeum equitans* and the recently cultured “*Nanopusillus acidilobi*” are the only ultra-small archaea that have been cultivated and characterized under laboratory conditions so far. The genome of *N. equitans* is only ~0.5 Mbp in size. It lacks nearly all genes essential for primary biosynthesis as well as the central and energy metabolism^14^, which indicates a strong dependency on its host *Ignicoccus hospitalis*. Interestingly, *I. hospitalis* cells do not divide if more than two *N. equitans* cells infect them. Still, generation time and final cell densities of *I. hospitalis* are not affected by infection^15^. Despite the absence of detectable damage or genomic regulatory changes to its host, *N. equitans* is described as a nutritional parasite^16^. Like *N. equitans*, “*Nanopusillus acidilobi”* is an obligate ecto-symbiont or parasite with a reduced genome (~0.6 kbp). It lacks the majority of the necessary genetic information for the energy metabolism and primary biosynthetic pathways except for several genes for proteins putatively involved in glycolysis/glyconeogenesis^17^.

The genomes of members of the not yet isolated Archaeal Richmond Mine Acidophilic Nanoorganisms (ARMAN) are ~0.5 Mbp larger than the genomes of *N. equitans* and “*N. acidilobi*”. They contain the necessary information for key functions of a central carbon metabolism^18^. This is indicative of a more independent lifestyle. The ARMAN were first detected by Baker *et al.* in 2006^19^. Since then, five different ARMAN groups (ARMAN-1 to ARMAN-5) were found in acidophilic biofilms that thrive at the Richmond Mine localized at the Iron Mountains in California^18^. The organisms are ellipsoid cells with volumes of 0.009 µm³ to 0.04 µm³ surrounded by a cell wall with only 92 ribosomes. Furthermore, most cells have enigmatic tubular structures of unknown function^20^. Genomic and proteomic data assembled from metagenomics data are available for ARMAN-2, -4 and -5. All three ARMAN have complete or nearly complete tricarboxylic acid cycles, but only ARMAN-4 and -5 have further genes for glycolytic pathways, the pentose phosphate pathway, and glycerol utilization^18^.

Still, the overall metabolism of the organisms, as well as the question of a putative dependency on other community members remain unclear. Moreover, prediction of the physiological capabilities of the ARMAN is highly complex if not even momentarily impossible since 25-38% of the predicted genes do not match sequences in public databases. Nevertheless, ARMAN are prominent members of submerged acid mine drainage (AMD) biofilms especially in anoxic parts^21^, which indicates a prominent role within these biocoenoses.

The 16S rRNA genes of ARMAN were also found in other ecosystems affected by AMD^22,23^. One of these is the former pyrite mine “Drei Kronen und Ehrt” in the Harz Mountains in Germany. Here, the microorganisms are embedded in gel-like biofilms, which resemble stalactite-formations^24^. These biofilms grow at average pH values of 2.0 to 2.5 and are characterized by an outer oxic shell and an inner anoxic core. The rim of the biofilm is mostly composed of chemolithoautotrophic bacteria that thrive using ferrous iron as energy and electron source and oxygen as electron acceptor. The anoxic center contains several different acidophilic archaea including ARMAN^25^.

This study describes the enrichment of ARMAN from biofilms of the “Drei Kronen und Ehrt” mine and the physiological, metagenomic and metatranscriptomic analysis of the enrichment culture. With long-term selective culturing experiments, it was possible to obtain very low diversity enrichment cultures of ARMAN related organisms. These are dominated by a not-yet cultured member of *Thermoplasmatales* and an ARMAN-1 organism. Analyses of the enrichment cultures as well as metagenomic and metatranscriptomic data offer new insights into the lifestyle of the previously uncultured ARMAN as well as two members of new families within the *Thermoplasmatales*.

## Results and Discussion

### Enrichment process

In 2013, Ziegler *et al.* described stalactite like biofilms that grow on the ceiling of an abundant pyrite mine in the Harz mountains in Germany^25^. Organisms belonging to the ARMAN as well as members of the *Thermoplasmatales* were detected within the anoxic core of the biofilms. The goal of this study was to produce a pure or at least highly enriched ARMAN culture in order to understand potential ecological dependencies of the organisms. Hence, the enrichment process used an anoxic medium containing casein and yeast extract as surrogates for biomass formed by chemolithoautotrophic primary producers of the biofilms. Furthermore, ferric sulfate was added as potential electron acceptor because iron and sulfate were prominent members of the water dripping from the biofilms^24^. Finally, the headspace of the hungate tubes was flushed with a hydrogen/carbon dioxide mixture (80%/20%) because there was also evidence for potential methanogenic activity in the biofilm, which could be supported by the release of hydrogen from fermentative microorganisms. A mixture of antibiotics was added to select for archaeal growth. After 1.5 years, bacteria could no longer be observed using CARD-FISH or PCR-analysis. Hence, the addition of antibiotics was omitted from this point onward.

After 2.5 years, the cultures showed a rather stable behavior and could be transferred every 10 weeks. Ferric iron reduction could be observed using the ferrozine assay within the individual growth intervals of the cultures. Moreover, addition of ferric sulfate was necessary for archaeal growth. Therefore, dissimilatory iron reduction seems to be an important trait of at least one member of the archaeal community. At this point, CARD-FISH analysis revealed the presence of archaea, ARMAN and a fungus (Supplementary Figure S1). Of note, the archaea probe does not hybridize to ARMAN species, which indicates the presence of at least one other archaeal species in the enrichment. Furthermore, preliminary PCR experiments targeting the 16S rRNA genes of archaea revealed the presence of at least one member of the *Thermoplasmatales* (data not shown).

### Metagenome and phylogenetic analysis

We could reconstruct the genomes belonging to three different archaea. Unfortunately, we could not find clear evidence for DNA or RNA that could be related to the fungus in the enrichment cultures. Nevertheless, the data showed a community built up predominantly by a novel member of the *Thermoplasmatales*, with the most closely related organisms *Thermogymnomonas acidicola* (91.6% identity, 1206 bp) and *Cuniculiplasma divulgatum* (91.7% identity, 1206 bp) (Supplementary Table S1 shows an overview of most closely related sequences) and an organism belonging to the ARMAN group. Surprisingly, the ARMAN 16S rRNA gene sequence had a 99% identity to a sequence belonging to the ARMAN-1 group that was obtained from samples collected at the Iron Mountains in California (USA) by Baker *et al.* in 2006^19^. So far, there are no genomic data available for this ARMAN group. A third and minor represented genome was also detected (Figure 1).

**Figure 1.**
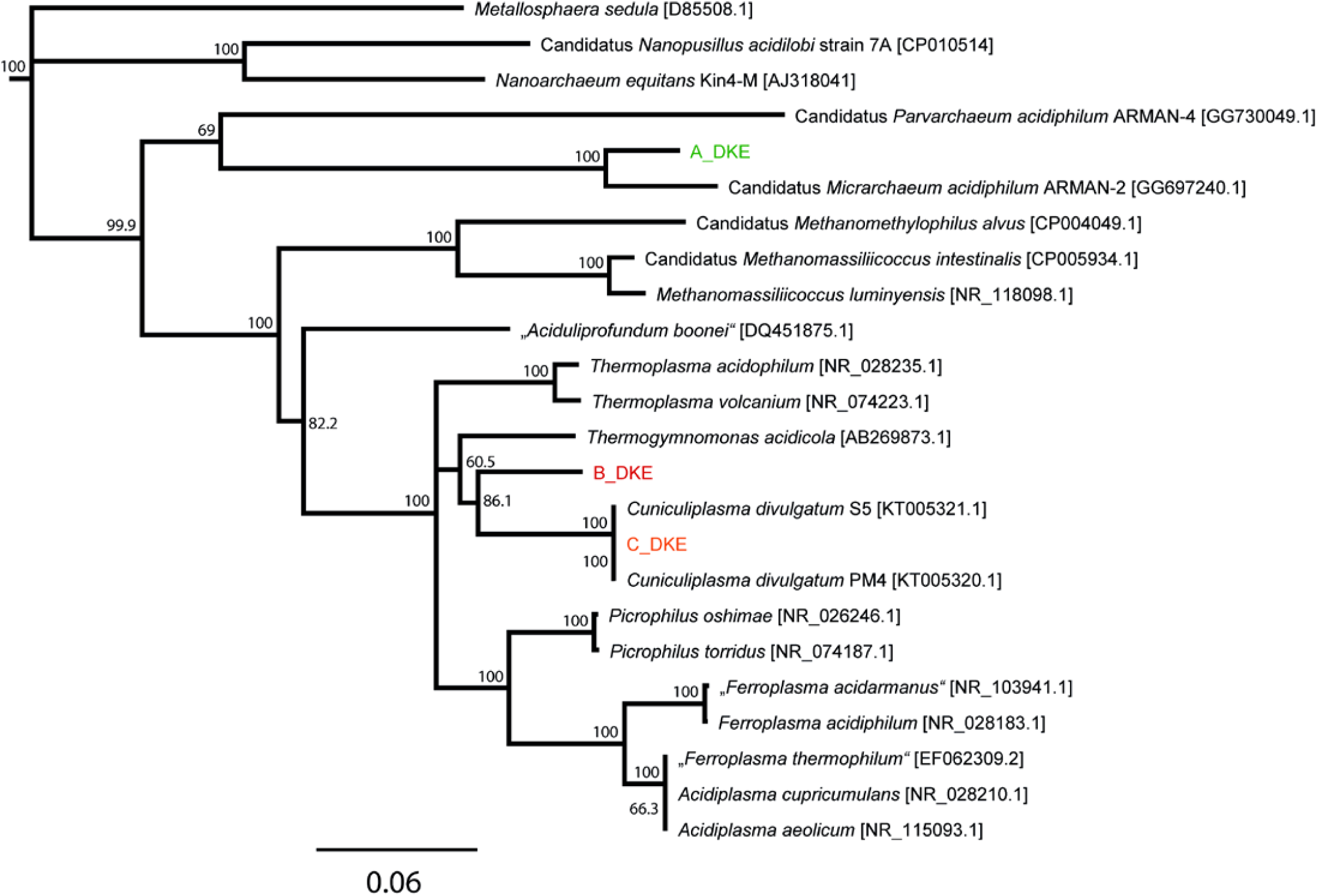
Phylogenetic tree of 16S rRNA gene sequences of all archaea found in the enrichment culture and related organisms. A data set based on the muscle alignment was used for tree building via the neighbor joining method. The bootstrap values were calculated from 1000 replicates, whereas values >50 are shown. The 16S rRNA gene sequence of ARMAN-1 was excluded, because it is incomplete and much shorter than all other published 16S rRNA gene sequences. An alignment with A_DKE shows a 99% identity. For 16S rRNA gene sequences of A_DKE, B_DKE and C_DKE see supplementary information.

The 16S rRNA gene sequence within this part of the metagenome showed a perfect match to the published sequence of the recently isolated *C. divulgatum*^26^ and to Gplasma, one member of the alphabet-plasmas^27^. Hence, it seems that three archaea and a fungus together form the stable consortium that was isolated over 2.5 years of successive transfers.

In the following sections, we will refer to the ARMAN-1 related organism as A_DKE (ARMAN_Drei_Kronen_und_Ehrt), to the novel member of the *Thermoplasmatales* as B_DKE and to the organism with the perfect match to the 16S rRNA gene sequence of *C. divulgatum* as C_DKE. We estimated the completeness for genome bins recovered from the metagenome samples and obtained values of 82% for A_DKE, 99% for B_DKE and 86% for C_DKE (Supplementary Table S2). Of note, analyses show that all available ARMAN-related genomes to date have a rather low predicted genome completeness by means of CheckM. This could indicate that ARMANs lack certain genes assumed to be present in all archaea.

The expected genome sizes of the three organisms are 1, 1.9 and 1.6 Mbp, respectively.

Using the genomic sequences, we sought to determine differences in the occurrences of gene families between A/B/C_DKE and also in comparison to available ARMAN sequences and genomes of five related members of the *Thermoplasmatales* (*Thermoplasma volcanium*, *T. acidophilum*, *Thermogymnomonas acidicola*, *Picrophilus torridus*, *Ferroplasma acidarmanus* and *Cuniculiplasma divulgatum* S5) (Figure 2). It is probably not surprising that there were significant differences in the overall number of gene families found in ARMAN genomes (~1009 +- 69 (SD)) compared to *Thermoplasmatales* genomes (~1,935 +- 557 (SD) (P < 0.004, two-sided Mann-Whitney U-test, Figure 2a). Significant differences were detected within a number of functional gene families annotated as translation, amino acid metabolism and metabolism of cofactor and vitamins among others (Figure 2b, Supplementary Table S3) (based on a two-sided Fisher's exact test, P=1.9E-8, 3.6E-8 and 5.0E-6, respectively). Within the four ARMAN genomes, we identified 227 gene families that together build the core genome. For the members of the *Thermoplasmatales*, the core genomes consist of 624 gene families. A dendrogram based upon the hierarchical clustering of gene families shared between genomes (Figure 2c) reveals that B_DKE and C_DKE cluster together with *T. acidicola*. A_DKE clusters as expected with the other available ARMAN genomes, whereas the closer relationship to ARMAN-2 (as seen by 16S rRNA gene analyses) is corroborated by the percentage of gene families shared between the ARMAN genomes.

**Figure 2.**
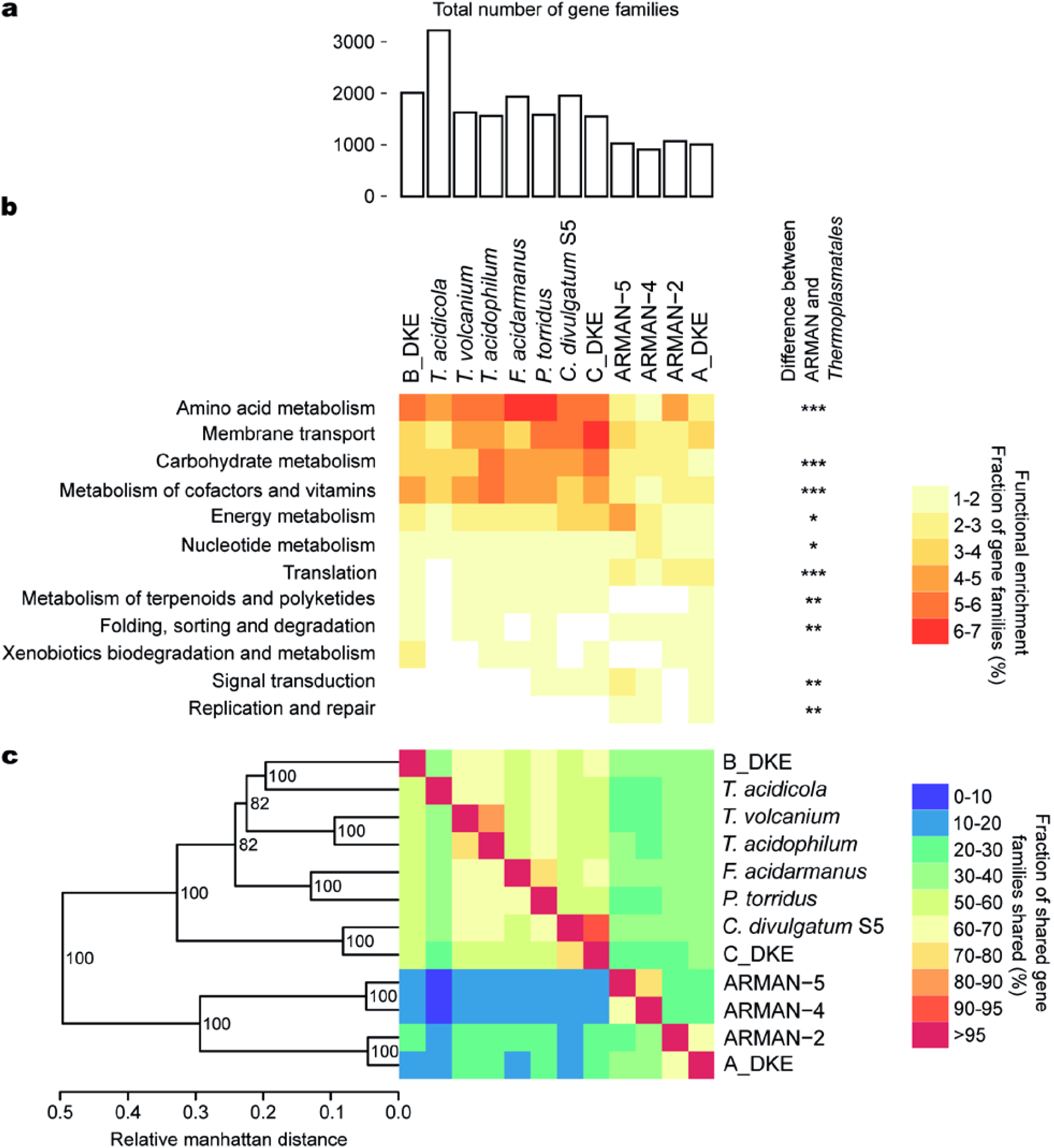
a) Total number of orthoMCL families found in indicated genomes. b) Functional enrichment analysis of orthologous genes. The heat map depicts the percentage of orthoMCL groups of each genome assigned to each indicated functional category. Stars indicate FDR corrected significance levels for difference in the number of orthoMCL families between the four ARMAN and eight *Thermoplasmatales* genomes (one star for value below ‘0.05’, two for ‘0.01’ and three for ‘0.001’. c) Heat map and hierarchical clustering dendrogram depicting the percentage of gene families shared between genomes. Node labels in the tree indicate bootstrap support after 100 iterations. Columns are normalized based on total number of gene families found in genomes.

A search for unique gene families (Figure 3c) in B_DKE compared to C_DKE revealed 914 unique gene families. Most of these were annotated as ABC transporters, Two-component system or as involved in the oxidative phosphorylation (more abundant in B_DKE) (Figure 3b). The A_DKE can be differentiated from the other sequenced ARMAN species by 227 unique gene families compared to 9 unique gene families shared by ARMAN-2, -4 and -5 (Figure 3c). We found that 15 of these 227 unique gene families could be successfully annotated (Supplementary Table S4). Compared to the other three ARMAN species, the genome of DKE_A shows more genes related to the benzoate degradation, phosphotransferase system, metabolism of aromatic compounds and amino acid metabolism and degradation (Figure 3a). Although the 16S rRNA gene sequence of C_DKE has an identical sequence to C. divulgatum, we could identify 80 unique gene families, which suggest that we have enriched a new strain of this species^28^.

**Figure 3.**
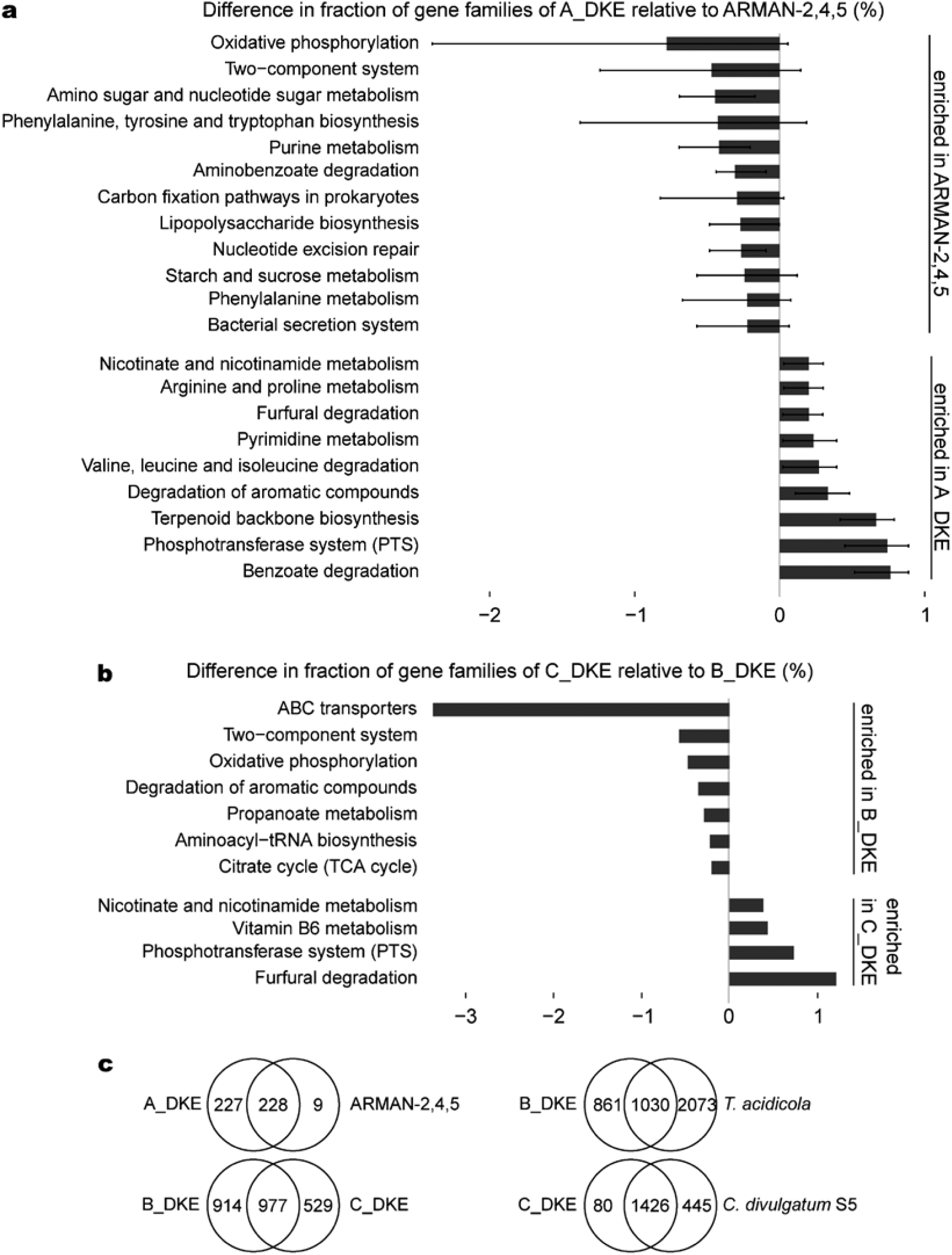
a) Bar plot showing the over-abundance of functions based on *de novo* inferred gene families based on orthoMCL analysis of three ARMAN genomes and the A_DKE genome. Length of the bars corresponds to the mean difference in the fraction of gene families annotated with indicated functional group. Lines show the difference to the ARMAN genomes with the minimal and the maximal fraction. Only functional groups are shown that differ in more than 0.2% or less than -0.2%. b) Bar plot showing the over-abundance of orthoMCL families assigned to indicated functional categories in C_DKE compared to B_DKE. Only functional groups are shown that differ in more than 0.2% or less than -0.2%. c) Venn diagram of gene families shared between indicated genomes. Gene families related to pathogenicity factors were not included in this overview.

### Metagenomic evidence for central metabolic properties

Co-enrichment of the three archaea and the fungus might be due to a metabolic dependency between the organisms. Therefore, we used the metagenomic information to screen for central metabolic capabilities, which would simultaneously highlight potential auxotrophic functions. The genomes of all three archaeal species contain several enzymes of the Entner Doudoroff (ED) pathway (Figure 4). Interestingly, the genome of B_DKE contains nearly all enzymes for the branched ED pathway including the key enzyme bifunctional KDG-/KDPG-aldolase. Nevertheless, we could not find a corresponding gene to a KDG-kinase (Supplementary Table S5). The genome of C_DKE shows all enzymes of the non-phosphorylated variant of the ED except its key enzyme the KDG-aldolase. Golyshina *et al.* (2016) stated that *Cuniculiplasma divulgatum* is incapable of growth on a number of sugar compounds, which is confirmed by our results^26^, whereas a complete non-phosphorylative ED was detected in the genome of G-plasma^27^. Furthermore, the activity of non-phosphorylative Entner Doudoroff pathways was experimentally shown previously in other related members of the *Thermoplasmatales* like *Thermoplasma acidophilum*^29^,*Picrophilus torridus*^30^ and *Cuniculiplasma divulgatum*^31^. Hence, this potential lack of pathways for the catabolism of sugar compounds is not a general characteristic of similar organisms.

**Figure 4.**
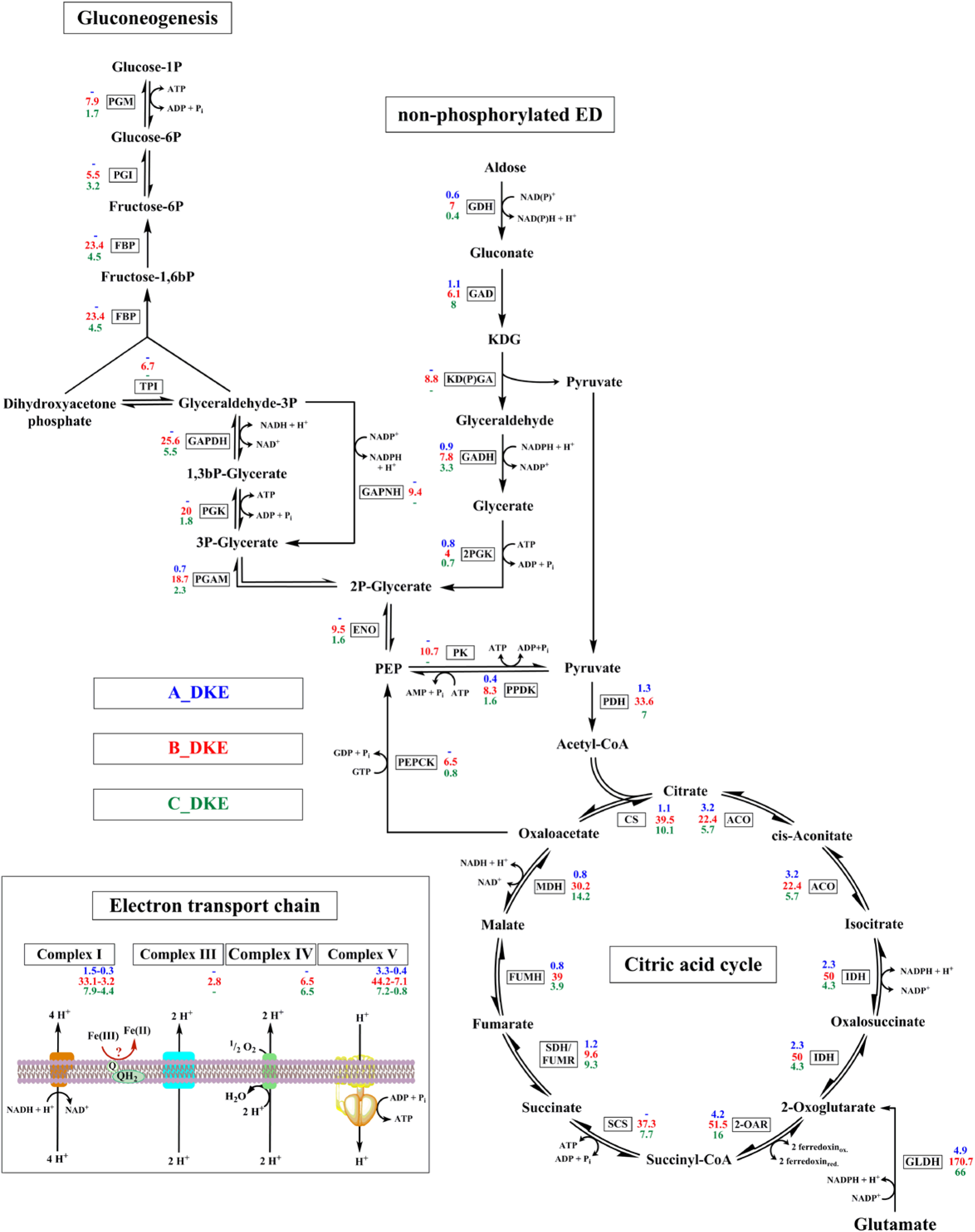
Postulated metabolism of all three archaea of the enrichment culture. Intermediates and catalyzing enzymes are indicated. The expression levels of the corresponding genes are listed. Blue numbers indicate RPKM-values of A_DKE, while red and green numbers refer to B_ and C_DKE expression, respectively. Abbreviations: PGM: phosphomannomutase/phosphoglucomutase, PGI: glucose/mannose-6-phosphate isomerase, FBP: fructose 1,6-bisphosphate aldolase/phosphatase, TPI: triosephosphate isomerase, GAPDH: glyceraldehyde-3-phosphate dehydrogenase (NAD(P)), PGK: phosphoglycerate kinase, PGAM: 2,3-bisphosphoglycerate-independent phosphoglycerate mutase, ENO: enolase, PEPCK: phosphoenolpyruvate carboxykinase (GTP), PPDK: pyruvate, orthophosphate dikinase; GDH: glucose/galactose 1-dehydrogenase (NADP+), GAD: gluconate/galactonate dehydratase, KD(P)GA: 2-dehydro-3-deoxy-D-gluconate/2-dehydro-3-deoxy-phosphogluconate aldolase, GADH: D-glyceraldehyde dehydrogenase, GAPN: glyceraldehyde-3-phosphate dehydrogenase [NAD(P)+], 2PGK: glycerate 2-kinase, PK: pyruvate kinase, PDH: pyruvate dehydrogenase; CS: citrate synthase, ACO: aconitase, IDH: isocitrate dehydrogenase, SCS: succinyl-CoA synthetase, SDH/FUMR: succinate dehydrogenase/fumarate reductase, FUMH: fumarate hydratase, MDH: malate dehydrogenase.

Sugar degradation catalyzed by the two *Thermoplasmatales* members via glycolysis can be excluded since both genomes lack evidence for an encoded phosphofructokinase, which corresponds well with the genomic data of *C. divulgatum*^31^. Still, B_DKE and C_DKE show all enzymes for gluconeogenesis, with the exception of the fructose 1,6-bisphosphate (FBP) aldolase. Nevertheless, a corresponding gene for this enzyme is missing in most archaeal genomes including *Thermoplasma acidophilum*^32^ and *Picrophilus torridus*^33^. Results by Say and Fuchs (2010) revealed the presence of a bifunctional FBP aldolase-phosphatase with high FBP aldolase and FBP phosphatase activity, which guarantees an unidirectional gluconeogenesis pathway^34^. This exists in nearly all archaeal groups and may be the putative ancestral gluconeogenic enzyme. A corresponding gene was found in the genomes of B_DKE and C_DKE, which likely allows both organisms to produce sugar compounds. The key enzyme for glycolysis, the phosphofructokinase, is missing also in the genomic data of Gplasma, whereas a bifunctional FBP aldolase-phosphatase is detectable^27^.

The genome of ARMAN-1 differs from previously sequenced ARMAN species in terms of sugar metabolism. Baker *et al.* (2010) described a pentose phosphate pathway for ARMAN-4 and -5 and an incomplete glycolysis for ARMAN-2^18^. In contrast, neither enzymes for glycolysis nor for gluconeogenesis or the pentose phosphate pathway were found in the ARMAN-1 genome. Moreover, the genes encoding for key reactions of the ED pathway were not detectable. Hence, there seems to be a necessity for the uptake of hexose and pentose sugars from the environment. In contrast, both *Thermoplasmatales* members contain all necessary enzymes for pentose formation via the non-oxidative pentose phosphate pathway similar to other members of *Thermoplasmatales*^35^.

All community members contain enzymes for a complete tricarboxylic acid cycle (TCA), with the exception that bioinformatic analysis based on the prokka database revealed so far only one subunit of the succinyl-CoA synthetase in A_DKE. In contrast, genomic analyses of *C. divulgatum* PM4 and S5 revealed that 2-oxoglutarate dehydrogenase, fumarate reductase and fumarase might be missing^31^. Nevertheless, Golyshina et al. point out that these enzyme functions could be functionally replaced by enzymes catalyzing closely related reactions that were predicted by bioinformatic analysis. In the here analyzed strains, the presence of succinate dehydrogenase/fumarate reductase and 2-oxoglutarate/2-oxoacid ferredoxin oxidoreductase are also evidence for the presence of a reverse TCA cycle, but ATP citrate lyase—the third indicator gene for the reverse TCA—is missing in all three genomes.

Neither one of the enriched archaeal species contain a complete fatty acid oxidation pathway as predicted by the KEGG database. While A_DKE does not seem to have any of the key functions for beta-oxidation, C_DKE and B_DKE lack one (beta-hydroxyacyl-CoA dehydrogenase) or two (beta-hydroxyacyl-CoA dehydratase and dehydrogenase) functions, respectively.

Archaeal electron transport complexes can have unusual compositions^36^. Nevertheless, bioinformatic analysis predicts the existence of a NADH-dehydrogenase, a respiratory succinate-dehydrogenase and an ATP-synthase in all organisms of the enrichment culture. Furthermore, B_DKE and C_DKE contain corresponding genes for cytochrome c-oxidase although genetic evidence for a cytochrome c protein is lacking. Both contain a Rieske FeS protein that is a characteristic component of complex III, and it forms a gene cluster in B_DKE with a cytochrome b6-like protein that could have complex III function^36^.

In addition, at least B_DKE has a gene which most probably encodes a copper protein belonging to the plastocyanin/azurin family, which could functionally replace cytochrome c^36^. We currently do not know how this electron transport core chain might be connected to ferric iron reduction, which is catalyzed at least by B_DKE within the community (see below). Furthermore, there is a gap between NADH-dehydrogenase and the cytochrome c oxidase in C_DKE, and it is unknown whether the latter might be a remnant of genomic reduction and selection towards a shorter electron transport chain involving only complex I and a cytochrome ubiquinol oxidase.

### Metatranscriptome

The ratio of sequence reads that can be referred to B_DKE, A_DKE and C_DKE is 1:0.5:0.27. Hence, all three archaea considerably contribute to the overall metatranscriptome of the study. The high content of RNA sequences that can be referred to A_DKE is surprising because all previous studies on ARMAN revealed a remarkably small number of ribosomes and because — as will be shown later — the number of ARMAN cells within the community is lower than the sum of B_DKE und C_DKE. We suggest that the considerably high content of A_DKE RNA within the metatranscriptome is evidence for active growth of the majority of ARMAN cells at sampling. The most highly expressed gene related to central carbon metabolism in B_DKE is similar to *gdh* coding for the glutamate dehydrogenases. Similarly, at least a *gdh*-like gene is also highly expressed in A_DKE and C_DKE. The encoded enzyme connects the catabolism of amino acids with the tricarboxylic acid cycle. In agreement with this, it can be observed that genes for TCA cycle enzymes are more highly expressed relative to the ED pathway. The question about the variant of the ED remains unclear, because B_DKE expresses the 2-dehydro-3-deoxy-(phospho)gluconate (KD(P)G) aldolase as well as all enzymes for the non-phosphorylated variant of the ED and the glyceraldehyde-3-phosphate dehydrogenase (GAPN) that is involved in the semi-phosphorylated variant of the pathway and could not be detected in other members of the *Thermoplasmatales*^35^. The C_DKE and B_DKE also express the complete set of genes necessary for gluconeogenesis and the pentose phosphate pathway. Furthermore, all detected genes encoding for electron transfer processes are highly expressed.

An analysis of the 20 most highly expressed genes (Supplementary Table S6) showed that both *Thermoplasmatales* species have high expression rates for oxidative stress proteins as well as proteins linked to electron transport process. Glutamate dehydrogenase and 2-oxoacid ferredoxin oxidoreductase are also highly expressed in C_DKE. In A_DKE, 14 of the 20 most highly expressed genes encode for hypothetical proteins. The most highly expressed protein in the ARMAN-1 transcriptome encodes for the translation initiation factor IF-2.

The detected high expression rates for oxidative stress proteins could be due to the presence of trace amounts of oxygen during the cultivation, which would also indicate that oxygen could be a potential electron acceptor for the organisms. Hence, growth experiments were conducted within a glove box containing a 95% N_2_/5% H_2_ atmosphere. If it all, we saw slower growth of the fungus within the enrichment culture. The archaea did not seem to be affected by the strictly anoxic conditions.

### Enrichment of B_DKE and isolation of the fungus

Interestingly, omitting the addition of antibiotics lead in some cultures to an increased growth rate of DKE_B. This resulted in cultures containing only DKE_B and the fungus. The fungus was isolated from these cultures using a solidified medium described by Baker et al. (2004)^37^. Fungal colonies on these plates were used for phylogentic analysis using a primer set for ITS amplification. Of note, growth of other colonies besides the fungus was not observed. The phylogenetic classification of the 5.8S rRNA gene and relating ITS-DNA sequences of the fungus revealed a 99% identity to *Acidothrix acidophila,* which is an acidophilic fungus first isolated from acidic soils in the Czech Republic ^38^. It was isolated from acidic kaolin quarries with a pH of 1.5-2.5 and exposed sulfur-rich brown coal beds. Of note, growth of the fungus in the medium used for the enrichment process did not lead to a reduction of the ferric sulfate. In contrary, growth of a culture containing only the fungus and DKE_B was connected to ferric sulfate reduction. Hence, at least this archaeon was capable of ferric iron reduction.

### Timeline experiments

The dynamics and interactions within the enrichment cultures were analyzed via timeline experiments. Hence, triplicate cultures were analyzed over eleven weeks of growth. The cultures were fixed for CARD-FISH analyses twice weekly, and DNA was isolated for qPCR-based quantitative community analyses once a week. Furthermore, samples were taken for measurements of ferric iron reduction as well as control of the pH values every week. Of note, as was emphasized before, we do not have genetic information regarding the fungus so far. Therefore, we could not conclude with certainty from gene to cell quantities. Hence, the qPCR analysis was conducted only with the archaeal members of the consortium.

Analysis of the qPCR and CARD-FISH data revealed a rather similar growth behavior of A_DKE as well as B_DKE and C_DKE (Figure 5). In the lag phase, the cell numbers increased continuously with a slightly higher amount of A_DKE cells compared to that of B_DKE and C_DKE. By week six, all organisms seemed to enter the logarithmic growth phase. From this point on, B_DKE and C_DKE began to dominate the cultures. The highest cell concentrations were reached in week eight. Thereafter, a rapid decrease in cell numbers was detected. The CARD-FISH pictures also highlight that A_DKE cells are mostly part of B_DKE or C_DKE cell agglomerates. Assuming that agglomerates are formed by extracellular polymeric material, which usually consists to a large fraction out of sugar compounds, we speculated that the biofilm growth might ensure the access to sugar for A_DKE. We emphasized at the beginning that some cultures showed faster growth during the conducted transfers and that these cultures were composed only of B_DKE and the fungus. These cultures do not show growth in the form of flocks but rather are uniform planktonic cells. Hence, the change in the growth phenotype might be the reason for decreasing A_DKE cell numbers. Supplementary Figure S2 displays the difference between the two growth phenotypes of B_DKE.

**Figure 5.**
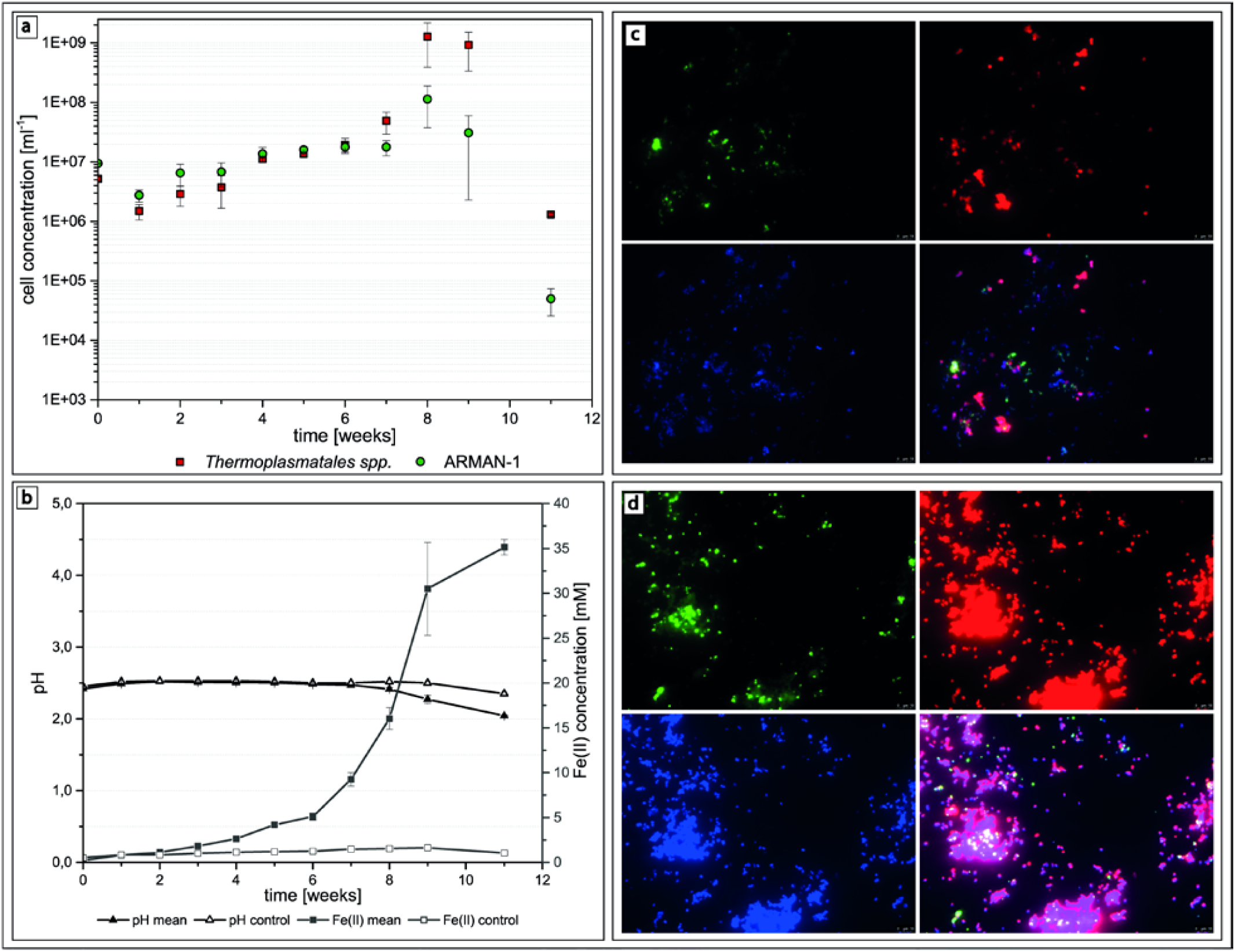
a) Cell concentrations of the enrichment cultures calculated from qPCR data of A_DKE, B_DKE and C_DKE, b) with corresponding pH values and Fe(II) concentrations over a time period of 11 weeks. Corresponding CARD-FISH pictures of enrichment cultures showing: c) a culture containing B_DKE and C_DKE (red, ARCH915, Alexa 546) and A_DKE (green, ARM980, Alexa 488) in agglomerates and d) an enrichment culture containing only B_DKE (red, ARCH915, Alexa 546), growing as single cells. Of note, the CARD-FISH fluorescence signal is not suitable to establish correct cell sizes and cells will most probably appear larger than they are.

Interestingly, the ferrous iron concentrations as well as the measured pH values correlate with the growth curve data (Figure 5). Until week six, only a slightly increased content of ferrous iron could be measured, and the pH values remained constant. From week seven on, there was a rapid ferric iron reduction along with increased growth rates. Even the pH values showed a slight decrease from this point on representing the acidification associated with the ferric iron reduction.

## Conclusion

We established a highly enriched consortium consisting of two acidophilic *Thermoplasmatales* species, an ARMAN, and a fungus. Of note, this is the only stable enrichment culture described so far containing ARMAN organisms. Moreover, it is the first report of an ARMAN-1 genome. The identified ARMAN-1 has a complete or nearly complete TCA cycle and completely lacks all enzymes necessary for glycolysis/glyconeogenesis and the pentose phosphate pathway. Moreover, we could not find evidence for gluconeogenesis and the pentose phosphate pathway, suggesting a dependency on synthesis of hexose and pentose sugars by other community members. This dependency seems to be corroborated by the identification of ARMAN cells within biofilm flocks that are likely built primarily by B_DKE. The colocalization of the cells might simplify access to suitable carbohydrates for the ARMAN. At least B_DKE seem to thrive using a respiratory metabolism with ferric iron as electron acceptor.

We tried to exclude oxygen from the growth medium and measurements of the oxygen content of the cultures by an optode suggest an anaerobic lifestyle. Moreover, similar growth of the archaea was also observed in an anoxic glove box. The reatively high expression levels of genes corresponding to enzymes involved in the elimination of oxidative stress might be due to the relatively short exposure to oxygen at the point of RNA sampling. In future experiments, we will combine heat inactivated B_DKE/fungus extracts with filtration based selection for pure ARMAN cultures. Nevertheless, B_DKE cells are also very small and pleomorphic, and the doubling time of the organisms is rather low. Hence, although future experiments seem straight forward they will require months or years of incubation time for several necessary culture transfers.

## Experimental Procedures

### Culturing conditions

Enrichment cultures of *Thermoplasma* spp. and ARMAN-1 originate from acidophilic stalactite-like biofilms of the former pyrite-mine “Drei Kronen und Ehrt” in the Harz Mountains, Germany (Ziegler et al. 2009 and 2013). Biofilms were inoculated under anoxic conditions in Hungate tubes and in a medium originally designed for the isolation of *Picrophilus* species ^39^. The pH of the medium was adjusted to 2.5 using 0.5 M H_2_SO_4_. Afterwards, the medium was autoclaved and supplemented with sterile filtered solutions of yeast extract, casein and ferric sulfate to final concentrations of 0.1% each and 20 mM, respectively, as critical additives. The growth of the bacterial species was repressed with 150 µg ml^-1^ streptomycin, 50 µg ml^-1^ kanamycin, 30 µg ml^-1^ chloramphenicol and 2 µg ml^-1^ vancomycin. The headspace was flushed with an 80% H_2_/CO_2_ 20% gas mixture. Cultures were inoculated with 10% of a good growing co-culture and incubated at 22°C. *Escherichia coli* cells were routinely cultured in LB medium supplemented with 40 µg ml^-1^ kanamycin or 100 µg ml^-1^ ampicillin, if necessary. Isolation of a fungus was conducted with a medium based on the study by Baker et al. (2004). The pH of the medium was adjusted to 2 using 0.5 M H_2_SO_4_ and it was supplemented with SL10 trace elements (DSMZ medium 320). The media plates were prepared with the addition of 2 % (w/v) agar.

### CARD-FISH

Samples were fixed for 1 h in 4% paraformaldehyde, washed twice in phosphate-buffered saline (PBS) and stored at -20°C in 50:50 PBS/ethanol. The fixed cells were hybridized and amplified as described previously ^25,40^. The following HRP-labelled 16S rDNA probes were used: ARCH915 (Archaea domain, GTG CTC CCC CGC CAA TTC CT, 20% formamide; Stahl and Amann, 1991) and ARM980 (ARMAN, GCC GTC GCT TCT GGT AAT, 30 % formamide; Baker *et al.*, 2006)). For amplification, Alexa_546_ and Alexa_488_ were used and counterstaining was conducted with DAPI. Samples were viewed on a Leica DM 5500B microscope (object lens 100x: HCX PL FLUOTAR, 1.4, oil immersion; eyepiece 10x: HC PLAN s (25) M), and images were taken with a Leica DFC 360 FX camera and the corresponding Leica LAS AF Lite software.

### Ferrous iron quantification

Ferric iron reduction was determined by quantification of the ferrous iron content spectrophotometrically using the ferrozine assay as described previously ^42^.

### Isolation of fungal DNA and amplification

A protocol based on Cenis (1992) was used to isolate DNA from the fungus. For taxonomic classification, the internal transcribed spacer (ITS) region primers ITS1 and ITS4 ^44^ were used to amplify the DNA with the iProof™ High-Fidelity PCR Kit (Bio-Rad, Munich, Germany).

### Quantification of cells using quantitative PCR

Quantitative PCR was used to quantify growth of the archaeal species in the enrichment culture. Standard curves were developed by integrating plasmids containing the target sequences of the qPCR primer sets in the genome of *E. coli*. Primer sets were developed for the 23S rRNA gene of ARMAN-1 and the 23S rRNA of the two *Thermoplasma* spp. using NCBI/Primer-BLAST ^45^ as described in the supplemental information. The primer sequences are listed in Table S7.

The organisms could contain more than one copy of the 23S rRNA gene, and this would hamper quantitative analysis. Thus, the normalized per-sample coverage of all genes was compared to the 23S rRNA gene converage. The latter was always below the overall coverage (Supplementary Table S8). Hence, the assumption was made that the 23S rRNA was represented by one copy in the genome.

### DNA/RNA isolation for Metagenomics/Metatranscriptomics

Isolation of genomic DNA for metagenomics analysis was conducted according to Lo et al. (2007). The total RNA isolation was conducted according to the protocol of the TRIzol Max Bacterial RNA Isolation Kit (Thermo Fisher, Waltham, USA) with 8 ml of an enrichment culture as the starting material. The remaining DNA was hydrolyzed with the DNA-*free*™ DNA Removal Kit (Thermo Fisher, Waltham, USA) following the manufacturer´s instructions.

### Metagenomic/metatranscriptomic sequencing and analyses

Metagenomic DNA from two analogous enrichment cultures was used for Illumina-based sequencing using a 2 x 51 bp protocol. The metatranscriptomic analysis was based on an RNA isolation from 8 mL of the enrichment culture. This sample was pooled from analogous enrichment cultures with similar growth characteristics. The RNA was sequenced using the 2 x 100 bp protocol. Supplementary Table S9 shows the overall statistics of the sequencing experiments.

### Accession codes

All DNA and RNA sequences that were retrieved within this study are publically available through NCBI BioProject: PRJNA358824.

## Acknowledgements

We gratefully thank Dr. Sibylle Bartsch and Dr. Katharina Geiger for fruitful discussions and are grateful for funding by the DFG.

## Sequences

### A_DKE 16S rRNA gene

ATTCCAGTCGATGCTGCTGGAGGGCACTGCTATCAGATTTCGACAGCCATGCAAGTTCGCTCAGCGGCTTCGCTGGGCAGCGTACGGCTCAGTAACACGTAGTCAATCTACCGTAAAGAGGCGAATACCCATGGGAAACTGTGGCTAATGCGCCATAGGCCAGGGATTCTGGAAGGAGCCCTGACTTAAAGGAGCGCGAATTGCAGCGCTTCCGCTTTACGATGAGACTGCGGCGGATCAGGCAGATGGCGGGGTAACGGCCCACCATACCTATAACCCGTAGGGGATGTGGGAGCATAAGCCCCGAGAAGGGCACTGAGACAAGGGCCCTAGCACTACGGTGTGCAGCAGGCGCGCAAACTCCACAATGCGCGTAAGCGTGATGGGGGGAATCTGAGTGGTTCACTTTGTGAACCTTTTGCCAAGTATAGCAAGCTTGGCGAATAAGTGCTGGGCAAGACCGGTGGCAGCCGCCACGGTAATACCGGCAGCACAAGTGGTGTCCACGATTATTGGGCTTAAAGCGCTCGTAGCCGGCTGATGCCGTCTCGCGTGAAATATTGGCGCTCAACGTCAATGCGTGCGCGAGATACCCATCGGCTAGGGAGCGGGTGAGGTCAGGAGTACTTATGGGGGAGGGGTTAAATCCTGTAATCCTATAAGGACTAACGGTGGCGAAGGCGCCTGACTAGAACGCATCCGACGGTGAGGAGCGAAGGCTAGGAGAACGAATCGGATCAGATACCCGAGTAGCCCTAGCAGTAAATTATGCAGACTCTGGTGTTGCAAGTATCACGAATGCTTGCAGTACCGTAGCGTAAGTGTTAAGTCTGCCGCCTGGGGAGTACGGCCGCAAGGTTGAAACTTAAAGCAATTGGCGGGGGAGCACACAAGGGGTGGATGCTGCGGTTTAATTGAATCCAACGTCGGAAATATTACCAGAAGCGACGGCAGTGTGAAGGTCAGACTGAAGATCTTACCTGACAAGCCGAGAGGTAGTGCATGGCGACCGTCAGCTCGTGGTGTGAACTGTCCGGTTAAGTCCGGTAAGTCTATGAGTGATACAATGACACAAAAGCTAAGACTAACGCCTGATACAAGCTACATGCTTGGAATATACAGGTGCAACAGGGGAAAGCGGATTGAACTGTCCTCAAGCGACGATGATATGATTGCAAGGTTTGCCAAGCTCGCAATGGACGAGTTCGGCATTGAGCCTAACAAGATAATCATCGACAGCGAGGGCAAGGTCAAGAAAACTTTCTTCTACAACTCCACGATGGTGAAAAGGTTTGAAAAGGCGCTTGAAAGAAGAGTAAACATATTCAAGTACGCCAATGCATATTCCGGAAATTACTTCGCGGCGCTGTTCGACTGCAACGGAGGCAAGGATGCTAAAGGCATCTTTCTGAAGGGCATGGACAGCGTTGACGAGGTAGTACTCGAAAGGCTCAACATACACACGGACAAAAGGGGCAGCAAGGATTACCTTGCAAACCAGTCCGCATTCATAACGCTAATAAAAGATTACAGCCTTAGAATTGCGAGTATTATTCATTGACCCGGAAACGAGCGAGACCCTCGCTTACAGTTGCTACCTGCTTAGAGATAAGCAGGGCACTCTGTTAGGACCGCCTTCGTTTAAGGAGGAGGAAGGAGAGGGCGACGATACGTCAGTATGCTCTGAATCTTCTGGGCTACACGCGGCATACAAAGGTCGGCACAATGAGATGCAACACCGAAAGGTGAAGCTAATCCCCTAAAACCGGCCCAAGTTCGGATTGAGGGCTGAAACTCGCCCTCATGAAGCTGGAATCCCTAGTAATCGCTTGTCACTATCGAGCGGTGAATACGTCCCTGCTCCTTGCACACACCGCCTACCAAGCCACCCAAGTGAGGTCTTAGTGAGGCAGCGCCACTGGCGTTTTCGAACTAAGGTTTCGCGAGGTGGGCTAAGGTATAACAAGGTATCCGTAGGGGAACCTGCGGATAGATCACCTCAA

### B_DKE 16S rRNA gene

GACCCCGGCGTACCGCTCAGTAACACGCGGATAATCTACCCCCAGGTGAGGTATAACCTCGGGAAACTGAGGATAATCCCTCATAGTTATCATATGCTGGAATGCTTTGATGACGAAAGCCTTTACGCCTGAGGATGAGTCTGCGGCCTATCAGGTCGTATGTGATGTAAAGGACCACATAGCCTAGGACGGGTACGGGCCCTGAAAGGGGGAGCCCGGAGATGGACTCTGAGACACAAGTCCAGGCCCTACGGGGCGCAGCAGGCGCGAAAACTGTGCAATGCGCGAAAGCGCGACACGGGAAACCTGAGTGCCTTGACAATGTCAAGGCTTTTCTTAAGCCTAAAAAGCTTAAGGAATAAGAGCTGGGCAAGACGGGTGCCAGCCGCCGCGGTAACACCCGCAGCTCGAGTGGTGGTCACTTTTATTGAGCCTAAAGCGTTTGTAGCCGGTTTTGCAAATCTTCAGATAAATTCCTCTGCTTAACAGATGATCTTCTGAAGAGACTGCGAGACTTGGGACCGGGTGAGGTTGAAGGTACTTCCGGGGTAGGGGTAAAATCCTGTAATCCTGGAAGGACGACCGGTGGCGAAGGCGTTCAACTAGAACGGATCCGACGGTGAGGAACGAGGGCTAGGGTAGCAAACCGGATTAGATACCCGGGTAGTCCTAGCTGTAAACACTGCCCACTTGGTGTTGCCCCTCCGATGAGGGGAGGCAGTGCCGGAGCGAAGGTGTTAAGTGGGCCGCTTGGGGAGTATGGTCGCAAGGCTGAAACTTAAAGGAATTGGCGGGGGAGCACCGCAACGGGAGGAGCGTGCGGTTTAATTGGATTCAACGCCGGAAAACTCACCGGGAGCGACCTTTGGATGAGAGTCAGCCTGATGAGCTTACTCGATAGAAGGAGAGGTGGTGCATGGCCGTCGTCAGCTCGTACCGTAGGGCGTTCACTTAAGTGTGATAACGAGCGAGACCCCCATCTCTAATTGCTAACGTGCATTCGCGTGCACGCGCACTTTAGAGGGACCGCCAGTGCTAAACTGGAGGAAGGAGGGGTCGACGGCAGGTCAGTACGCCCCGAATCTCCCGGGCAACACGCGCGCTACAAAGGGCAGGACAATGAGTTGCCACCTCGAAAGGGGGAGCTAATCTCGAAACCTGTTCGTAGTTAGGATTGAGGGCTGTAACTCGCCCTCATGAATGTGA

### C_DKE 16S rRNA gene

ACTCCGGTTGATCCTGCCGGCGGCTACTGCTATCAGGTTTCGACTAAGCCATGCGAGTCAAGGGGTCGTAAGACACCGGCGTACTGCTCAGTAACACGCGGACAATCTACCCCCAGGTGGGGGATAACCTCGGGAAACTGAGGCTAATACCCCATAGTTGTCATGTGCTGGAATGCTTTGACGATGAAAGAATTTCGCCTGAGGATGAGTCTGCGGCCTATCAGGTAGTATGTGGTGTAAAGGACCACATAGCCTAAGACGGGTACGGGCCTTGAAAGAGGGAGCCCGGAGATGGACTCTGAGACACTAGTCCAGGCCCTACGGGGCGCAGCAGGCGCGAAAACTGTGCAATGCGCGCAAGCGCGACACGGGAAGCCTGAGTGCCTTAACTATGTTAAGGCTTTTCTTGATCCTAAAAAGCTCAAGGAATAAGAGCTGGGTAAGACGGGTGCCAGCCGCCGCGGTAACACCCGCGGCTCGAGTGGTAGTCACTTTTATTGAGCCTAAAGCGTTTGTAGCCGGATTTGTAAATCTCTAGGAAAATACTTCTGCTTAACTGAAGAAATTTTGGAGAGACTGCAAATCTAGGGACCGGGTGAGGTTGAATGTACTTCTGGGGTAGGGGTAAAATCCTGTAATCCTGGAAGGACGACCGGTGGCGAAGGCGTTCAACTAGAACGGATCCGACGGTGAGGAACGAAGGCTAGGGGAGCAAACCGGATTAGATACCCGGGTAGTCCTAGCTGTAAACACTGCCCACTTGGTGTTGCATCTCCGGTAAGGGGGTGCAGTGCCGTAGCGAAGGTGTTAAGTGGGCCGCTTGGGGAGTATGGTCGCAAGGCTGAAACTTAAAGGAATTGGCGGGGGAGCACCGCAACGGGAGGAGCGTGCGGTTTAATTGGATTCAACGCCGGAAAACTCACCGGAGGCGACCTGTGTATGAGAGTCAACCTGATGAATTTACTCGATAGCAGGAGAGGTGGTGCATGGCCGTCGTCAGCTCGTACCGTAGGGCGTTCACTTAAGTGTGATAACGAGCGAGACCCCCATCTTCAATTGCTAAGCTAACCGTGAGGTTGGTAGAACTTTGAAGGGACCGCCAGTGCAGAACTGGAGGAAGGAGGGGTCGACGGCAGGTCAGTACGCCCCGAATCTTCCGGGCAACACGCGCGCTACAAAGGACAGGACAATGGGCTGCGACCTCGAAAGGGGAAGCCAATCCCGAAACCTGTTCGTAGTTAGGATTGAGGGCTGTAACTCGCCCTCATGAATCTGGATTCCGTAGTAATCGCGGGTCAACATCCCGCGGTGAATATGCCCCTGCTCCTTGCACACACCGCCTGTCAAACCATCCGAGCTGGTGTTGGATGAGGGTTAGTTCGAAAGGGTTGATTCGAATCTGATGTCAGTGAGGAGGGTTAAGTCATAACAAGGTATCCGTAGGGGAACCTGCGGATGGATCACCTCCT

### Fungus ITS

GGCGCGGGGGGTGCCGCCGGCGGCCAGCTCAACCCTCAACATTTTGAACCTGAGTGTCACAAATATAATAACCAAATTACAACCTGCAACAATGGATCTCTTGGTTCCGGCATCGATGAAGAACGCAGCGAAATGCGATACGTAATGTGAATTGCAGATCTCCAGTGAATCATCGAATCTTTGAACGCACATAGCGCCCGCCAGCACTCTGGCGGGCATGCCCGTCCGAGCGTCGATGACGCCGCTCGAGCCCGCCCTCCCCCTGTGGGGAGGAGGGCTCGGTGTTGGGGCGCTGGGGGTAGAACCCCAGGCCCCCAAAAGCATTGCGGGGTCCCCGCGCGGACC

